# Inhibition of granuloma triglyceride synthesis imparts control of *Mycobacterium tuberculosis* through curtailed inflammatory responses

**DOI:** 10.1101/2021.05.10.443218

**Authors:** Stanzin Dawa, Dilip Menon, Prabhakar Arumugam, Akash Kumar Bhaskar, Moumita Mondal, Vivek Rao, Sheetal Gandotra

**Affiliations:** Academy of Scientific and Innovative Research (AcSIR), Ghaziabad- 201002, India; Cardiorespiratory Disease Biology, CSIR-Institute of Genomics and Integrative Biology, New Delhi, India

**Author notes:** to whom correspondence must be addressed.

## Abstract

Lipid metabolism plays a complex and dynamic role in host-pathogen interaction during *Mycobacterium tuberculosis* infection. While bacterial lipid metabolism is key to the success of the pathogen, the host also offers a lipid rich environment in the form of necrotic caseous granulomas, making this association beneficial for the pathogen. Accumulation of the neutral lipid triglyceride, as lipid droplets at the cellular cuff of necrotic granulomas, is a peculiar feature of pulmonary tuberculosis. The role of triglyceride synthesis in the TB granuloma and its impact on the disease outcome has not been studied in detail. Here, we identified diacylglycerol O-acyltransferase 1 (DGAT1) to be essential for accumulation of triglyceride in necrotic TB granulomas using the C3HeB/FeJ murine model of infection. Treatment of infected mice with a pharmacological inhibitor of DGAT1 (T863) led to reduction in granuloma triglyceride levels and bacterial burden. A decrease in bacterial burden was associated with reduced neutrophil infiltration and degranulation, and a reduction in several pro-inflammatory cytokines including IL1β, TNFα, IL6, and IFNβ. Triglyceride lowering impacted eicosanoid production through both metabolic re-routing and via transcriptional control. Our data suggests that manipulation of lipid droplet homeostasis may offer a means for host directed therapy in Tuberculosis.

## Introduction

*Mycobacterium tuberculosis* (Mtb), the causative agent for Tuberculosis in humans, is the leading cause of mortality due to a single infectious agent. Even though treatment exists, infected individuals are at a life-long risk of poor lung function in case of pulmonary TB and debilitating morbidity in case of extrapulmonary TB (Byrne et al., 2015). Inflammation plays a significant role in these outcomes and thus host directed therapies that target inflammation and promote mycobactericidal activities of the host are sought in new treatment regimens (Young et al., 2020). Understanding the molecular events underpinning ensuing inflammation in the TB granuloma is key towards moving in this direction.

Granulomas with necrotizing centers are hallmarks of tuberculosis in humans. Neutral lipid laden macrophages, also termed foamy macrophages, form a cellular cuff at the rim of these necrotic centers (Jaisinghani et al., 2018; Kim et al., 2010; Peyron et al., 2008). Intracellularly, the neutral lipid is packaged in the form of lipid droplets that can be detected as oil red O positive droplets or as hematoxylin and eosin negative vacuoles that appear as “holes within a foam”, thus terming these macrophages as “foamy macrophages”. The origin of foamy macrophages is attributed to the ensuing inflammatory response, with roles for interferon gamma (IFNγ), tumor necrosis factor alpha (TNFα), lipids in the pleural effusion and necrotic milieu being identified as relevant players across host species (Genoula et al., 2018; Guerrini et al., 2018; Jaisinghani et al., 2018; Knight et al., 2018). Necrotic TB granulomas are distinctive from other atherogenic pathologies, in that both triglyceride and cholesterol ester accumulation takes place in TB while cholesterol esters are more abundant in atherogenic lesions (Guerrini et al., 2018; Kim et al., 2010). Proteomic characteristics of lipid droplets that are rich in triglycerides are distinct from those rich in cholesterol ester (Hsieh et al., 2012; Khor et al., 2014); this points towards distinct roles and interactions executed by these storage organelles. As the lipid droplet proteome of macrophages is dynamically altered upon *M. tuberculosis* infection, metabolism of these lipid classes in the pathophysiology of the TB granuloma may play different roles (Menon et al., 2019).

Macrophages are key hosts that are manipulated by *Mycobacterium tuberculosis* in the course of disease. Metabolic routes selected by a macrophage define the defence trajectory upon infection. Glycolytic flux polarizes macrophages towards the inflammatory M1 polarized macrophages. Interestingly, lung interstitial macrophages exhibit this phenotype and are capable of Mtb control (Huang et al., 2018). In contrast, resident alveolar macrophages exhibit commitment towards fatty acid oxidation and oxidative phosphorylation, and harbour higher bacterial burden (Huang et al., 2018). The handling of fatty acids towards storage visa-vis oxidation is another determinant of inflammatory state gained by cells. Tuberculous pleural effusion treated macrophages also induce cholesterol ester biogenesis, in a process dependent on IL10 driven induction of the enzyme acyl CoA:cholesterol acyl transferase (ACAT) (Genoula et al., 2018). This process renders the macrophage M2 polarized and promotes bacterial growth; the role of ACAT activity in control of infection remains to be addressed in this model. M1 polarization by IFNγ treatment also leads to accumulation of cholesterol esters via HIF1α activation (Knight et al., 2018; O’Neill and Pearce, 2016). This is attributed to the induction of the lipid droplet localized protein Hig2 (Knight et al., 2018). Myeloid specific deletion of HIF1α led to impaired control of mycobacteria in lungs of infected mice and also decreased accumulation of neutral lipids within granuloma macrophages (Braverman et al., 2016; Knight et al., 2018). However, Hig2 deficiency impacted only the neutral lipid content and not bacterial control (Knight et al., 2018). These studies have thus been able to separate the mycobactericidal capacity from lipid storage capacity in metabolically polarized macrophages.

Previous work has demonstrated that human macrophages treated with necrotic macrophages not only makes them triglyceride loaded, it also renders them hyper-inflammatory to Mtb infection (Jaisinghani et al., 2018). The final step in triglyceride synthesis is catalysed by the enzyme diacylglycerol-O-acyltransferase (DGAT1). Reduction in triglyceride levels by silencing of DGAT1 in human macrophages led to reduction of the inflammatory response to Mtb infection (Jaisinghani et al., 2018). More recently, LPS stimulated DGAT1 activity was also found to be important for the inflammatory response of murine macrophages (Castoldi et al., 2020). Triglycerides are a potential source of arachidonic acids, and macrophage lipid droplets harbour lipases and cyclo-oxygenases which can catalyze stepwise conversion of this esterified arachidonic acid to eicosanoids (Dvorak et al., 1992; Menon et al., 2019). Eicosanoids are important lipid mediators of inflammation which balance the amplitude of inflammatory response and mycobacterial control (Tobin et al., 2012). Even in the case of Mtb infection of murine bone marrow derived macrophages, triglyceride synthesis by DGAT1 contributes to the production of several eicosanoids (Knight et al., 2018). However, the control of infection in macrophages is independent of triglyceride synthesis (Jaisinghani et al., 2018; Knight et al., 2018).

The peculiar placement of triglyceride rich foamy macrophages at the interface of the cellular and necrotic region of the human TB granuloma led us to question the role of triglyceride synthesis in an *in vivo* model that mimics this pathology. Here we use the Kramnik mouse model (C3HeB/FeJ) wherein infection culminates in pulmonary necrotic lesions upon aerosol challenge with Mtb (Driver et al., 2012; Kramnik et al., 1998). A temporal onset of oil red O positivity in TB granulomas concomitant with macronecrosis offered a suitable model to pharmacologically target DGAT1. We demonstrate that DGAT1 inhibition prevents neutral lipid accumulation in necrotic granulomas and leads to improved control on bacterial burden. Triglyceride lowering mediated bacterial control was associated with reduction in neutrophil infiltration, expression of pro-inflammatory pathways and change in eicosanoid flux. Our study offers a fresh insight into the potential of targeting lipid synthesis as host directed therapy to restrict the inflammatory response and improve the host’s ability to control bacterial growth.

## Results

### High dose aerosol infection in C3HeB/FeJ mice exhibits progressive disease with caseous necrotic granulomas

Previous studies have revealed high heterogeneity in TB pathology in the C3HeB/FeJ mouse model, with only 50% of infected animals resulting in necrotic granulomas between 28 and 42 days post infection (Irwin et al., 2015). In addition, an inter-lobe variability in this model limits correlation between histology, inflammatory response, and bacterial burden. We infected C3HeB/FeJ mice with 100 or 500 cfu of Mtb Erdman via the aerosol route. Analysis of the left caudal lobe of mice revealed higher heterogeneity of mice infected with 100 cfu compared to 500 cfu. At d28 post infection with 100 bacilli, 9 out of 12 mice exhibited at least one necrotic lesion, 2 exhibited no necrotic lesions while one exhibited no lesion at all (FigS1A and B). In contrast, all mice infected with 500 bacilli exhibited at least one necrotic lesion in the left caudal lobe at d28 post infection. Given the reduced heterogeneity with the higher dose, we further characterized this high dose model of infection. Nodular lesions were not visible at d14 while airway thickening and cellular infiltration was visible (Fig 1A and B). Nodular macroscopic lesions were visible by d28 post infection in all infected animals. At this point, 60% of animals exhibited multiple large lesions and 40% exhibited coalesced lesions (Fig 1C). Mortality was evident starting from d35 and progression of disease was associated with visibly necrotic lesions.

**Figure 1.**
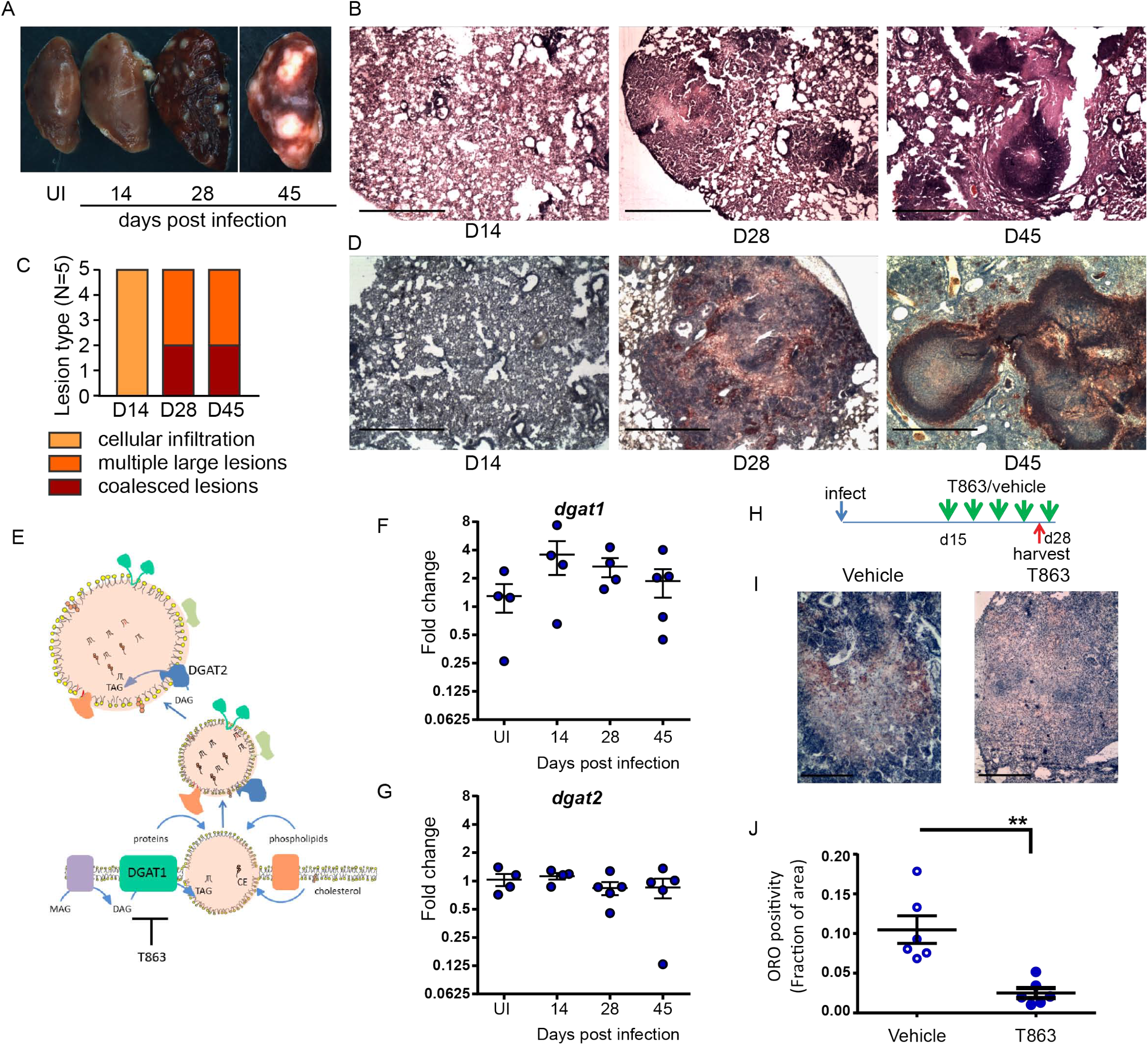
Neutral lipid accumulation in TB granulomas is Dgat1 dependent in susceptible mice. (A) Gross pathology of infected mice across time. (B) Representative images acquired at 2X magnification of hemotoxylin and eosin stained lung sections at the indicated time points. Scale bar= 1mm. (C) Quantitative assessment of lesion types post infection. (D) Oil red O and hematoxylin staining across indicated time points post infection. scale bar=1 mm. (E) Schematic representing stages of lipid droplet growth at which terminal enzymes involved in triglyceride synthesis are indicated. Expression of *dgat1* (F) and *dgat2* (G) post infection in lungs of C3HeB/FeJ mice.For panels 1-G, N=4-5 animals per time point. (H) Shematic representing intravenous treatment schedule with the DGAT1 inhibitor T863 and Vehicle indicaed with green arrows (once every three days). (I) Representative Oil red O and hematoxylin stained lung sections of vehicle and T863 treated mice at d28 post infection. Scale bar=400 μm. (J) ORO positivity, represented as fraction of oil red O positive regions normalized to total area of the left caudal lobe. Data are mean+/- SD from 6 animals per group with equal numbers of male and female mice per group. Horizontal bars in graphs represent mean+/- SEM. Statistical analysis: unpaired t-test with Welch’s correction; **p<0.01.

Human and guinea pig granulomas exhibit accumulation of neutral lipid loaded macrophages at the cellular cuff of necrotic granulomas (Jaisinghani et al., 2018; Peyron et al., 2008). We sought to determine if this aspect of human granulomas is reproduced in C3HeB/FeJ mice. We stained lung sections for the presence of neutral lipids using oil red O and found no staining at d14. By d28, the necrotic granulomas stained for oil red O, largely in regions that were devoid of lymphocytes (Fig 1D). By d45, well-formed caseous lesions with a central necrotic core were visible. Oil red O positive cells organized as a cuff surrounding this central necrotic core (Fig 1D).

### DGAT1 regulates granuloma triglyceride levels

Triglyceride (TG) is one of the major neutral lipids found in human and guinea pig granulomas (Jaisinghani et al., 2018; Kim et al., 2010). The synthesis of TG relies on either of two enzymes, DGAT1 or DGAT2. DGAT1 synthesizes TG at the ER membrane which is where the lipid droplet (LD) originates (Walther et al., 2017) (Fig 1E). DGAT2 also contributes to TG synthesis at the surface of the already formed LD. We next sought to understand whether these enzymes were induced upon infection. We observed 4-fold induction of *DGAT1* by d14 post infection (Fig 1F). Two fold Induction of *DGAT1* was maintained at d28, followed by greater animal to animal variation by d45 post infection. In contrast, *DGAT2* was not found to be induced upon infection at all (Fig 1G). The peak expression of *DGAT1* preceded the onset of neutral lipid accumulation, suggesting a role for DGAT1 in neutral lipid accumulation within the granuloma.

To understand if DGAT1 activity was essential for neutral lipid levels in the granuloma, we treated mice with a pharmacological inhibitor of DGAT1. T863 is a highly specific and potent inhibitor of DGAT1 previously found to be pharmacologically active (Cao et al., 2011). However, DGAT1 has also been found to be important for lipid clearance from distal regions of the small intestine (Lee et al., 2010). In C3HeB/FeJ mice treated orally with T863 at a dose of 30 mg/Kg over a period of 3 weeks, we found higher oil red O staining in the distal region of the small intestine (Fig S2A). Consistent with lipid malabsorption, we found a reduction in serum triglycerides in mice treated orally with T863 (FigS2B). To avoid this effect on dietary lipid clearance and enable delivery to the lung, we chose the intravenous route for delivery. Intravenous delivery via the tail vein of an equivalent dose was performed every three days over a period of 2 weeks, from d15 to d28 post infection, to monitor the effect on lung neutral lipid accumulation (Fig 1H). Intense oil red O positivity was observed in cells proximal to the central necrosis of granulomas in case of vehicle treated animals (Fig 1I). No such “cuff” of oil red O positivity was observed in T863 treated animals (Fig 1I). Quantification of the stain in whole lung sections revealed approximately 80% reduction in oil red O positivity upon T863 treatment (Fig 1J). These data revealed DGAT1 to be a major contributor to neutral lipid accumulation in TB granulomas. No significant difference was found in serum TG levels (Fig S2C). As *DGAT1* expression induced by d14, we further tested impact of treatment starting at d7 post infection. This treatment did not result in any further weight loss beyond that observed due to infection (Fig S3). Together, the intravenous dosing of T863 in a high dose model revealed DGAT1 activity to be essential for granuloma TG synthesis and was a suitable methodology to test the role of DGAT1 activity in infection progression.

### DGAT1 inhibition leads to control in pulmonary growth of *M. tuberculosis*

To measure the impact of granuloma triglyceride levels on disease progression, we estimated bacterial burden from lungs of vehicle and T863 treated animals by CFU plating at d28. The overall T863 group exhibited 40% reduction in mean bacterial burden while it was evident that T863 treated animals exhibited a dimorphic response in terms of pulmonary bacterial burden (Fig 2A). Five out of eleven animals exhibited 97-25% reduction while 6 out of 11 animals exhibited no change in bacterial burden compared to the vehicle group. To understand if the decreased bacterial numbers were due to a role of macrophage TG pools in providing essential nutrients to intracellular Mtb, we infected bone marrow derived macrophages from C3HeB/FeJ mice with Mtb and studied the impact of T863 on bacterial growth. We found no impact of T863 on intracellular bacterial growth (Fig 2B). Previously, IFNγ and IL6 have been shown to induce lipid droplet accumulation in murine bone marrow derived macrophages (Knight et al., 2018; Pean et al., 2017). To test if this polarization rendered bacterial control dependent on triglyceride levels, we polarized macrophages with each of these cytokines prior to infection. However, T863 did not impact control of Mtb across either of these polarization methods (Fig 2C and D). Recently, IL4 was shown to prevent lipid accumulation in an *in vitro* model of foamy macrophage formation (Genoula et al., 2020). In addition the IL13/IL4R axis is associated with aggravated pathology *in vivo* (Heitmann et al., 2014). We therefore also tested whether IL4 or IL13 (both M2 polarizing cytokines) could restrict Mtb growth in a triglyceride dependent manner. In neither of these conditions did we observe an effect of T863 on bacterial burden (Fig 2E, and data not shown for IL13). In addition, splenic bacterial burden was not affected by T863 treatment (Fig S4), suggesting a lung pathology specific restriction on bacterial growth by limiting TG synthesis.

**Figure 2.**
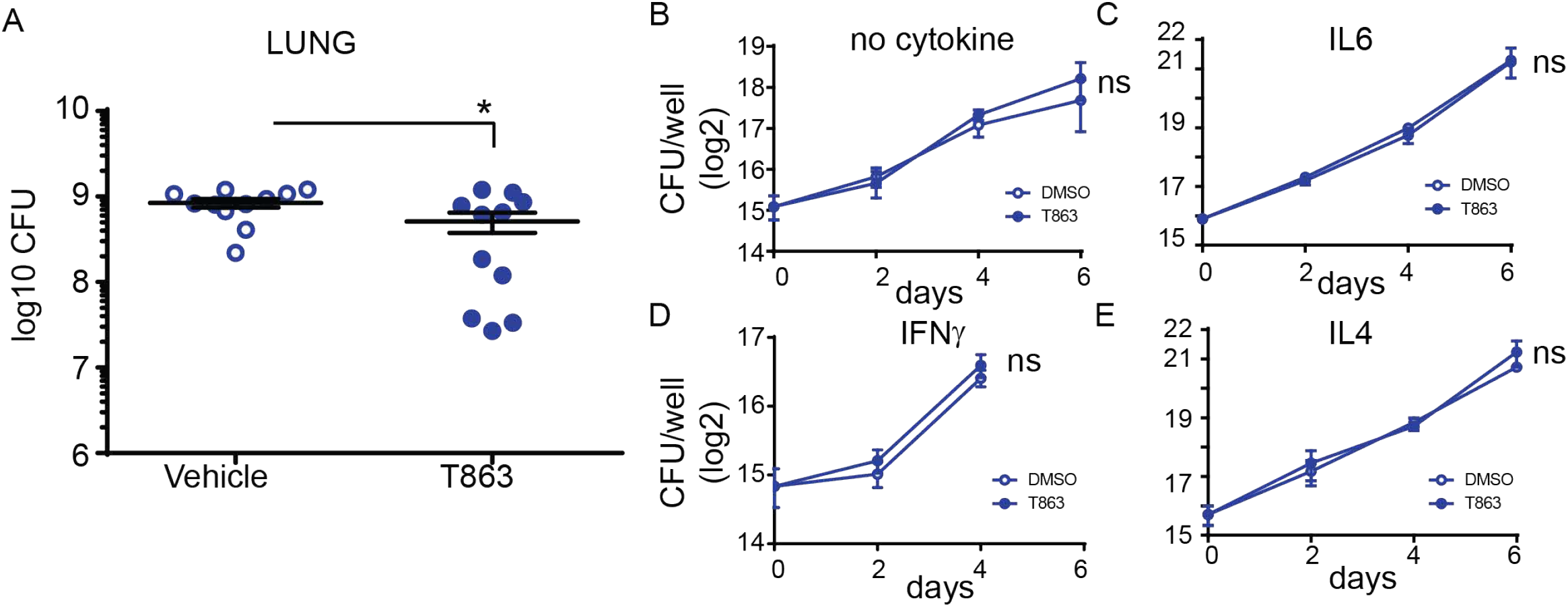
DGAT1 activity creates a growth permissive environment in the lung. (A) Lung bacterial burden in mice treated with T863 from d8 to d28. Horizontal bars in the graph represent mean+/- SEM from 11 animals/group. (B-E) Intramacrophage growth of Mtb upon T863 treatment under different stimuli for macrophage polarization. Data are mean+/- SEM from 2-3 independent experiments with triplicate infection wells in each experiment. Statistical analysis: Unpaired t-test with Welch’s correction, *p<0.05, ns=not significant.

### T863 mediated bacterial control correlates with reduced neutrophil infiltration

Given the heterogeneity in response to T863, we decided to investigate the molecular features that correlate with lowering bacterial numbers upon T863 treatment *in vivo*. Towards this, we performed total RNA-seq from lungs of both, vehicle and T863 treated animals. Given the inherent within-group heterogeneity, we found no differentially expressed genes between the vehicle versus T863 treated group (adjusted p-val<0.05, FC>2 or <0.5). Principal component analysis clustered the 6/11 animals that exhibited no change in cfu with the animals of the vehicle group and 5/11 animals that exhibited 97-25% reduction in bacterial burden away from these. We therefore defined two subgroups within the T863 treated animals: the low cfu group (T863^lo^) was defined as those that harboured <25% bacterial burden of the average cfu in the vehicle group, while the rest of the animals in the T863 treated group were defined as T863^hi^. A total of 2343 genes were found to be differentially expressed between the T863^hi^ and T863^lo^ groups while 1143 genes were found to be differentially expressed between Vehicle and T863^lo^ groups (Table S1 and S2). Only 5 genes were found to be differentially expressed between the T863^hi^ and Vehicle groups (increased >2 fold). These data suggest that albeit a global metabolic impact of T863 on local neutral lipid levels, impact on gene expression may be limited by inter-individual heterogeneity.

Pathway enrichment analysis of the differentially expressed genes revealed that neutrophil degranulation was the most significantly diminished pathway in the T863^lo^ group compared to both the vehicle and T863^hi^ group (Fig 3A and Table S3). Forty nine genes of this pathway were differentially abundant between T863^lo^ and Vehicle group and 66 genes were differentially abundant between T863^lo^ and T863^hi^ groups. A common subset of 24 neutrophil associated genes was reduced in expression in T863^lo^ group (Fig 3B). Neutrophils are known to release several matrix metallopeptidases (MMPs) that contribute to the granulomatous pathology. MMPs exhibit collagenolytic activity which is important for the initial stages of granuloma formation and for lung tissue destruction that is a hallmark of tuberculosis (Ong et al., 2014). Neutrophil specific Mmps (*Mmp8*, *Mmp9*, and *Mmp25*) were found to be significantly reduced in expression in T863^lo^ animals. In addition, expression of genes belonging to extracellular matrix organization and collagen degradation were also decreased in expression. Simultaneously, an increase in expression of genes belonging to muscle contraction and elastic fibre formation indicated the ability of T863 to promote restoration of a healthy lung architecture in the T863^lo^ group(Fig 3C and D).

**Figure 3.**
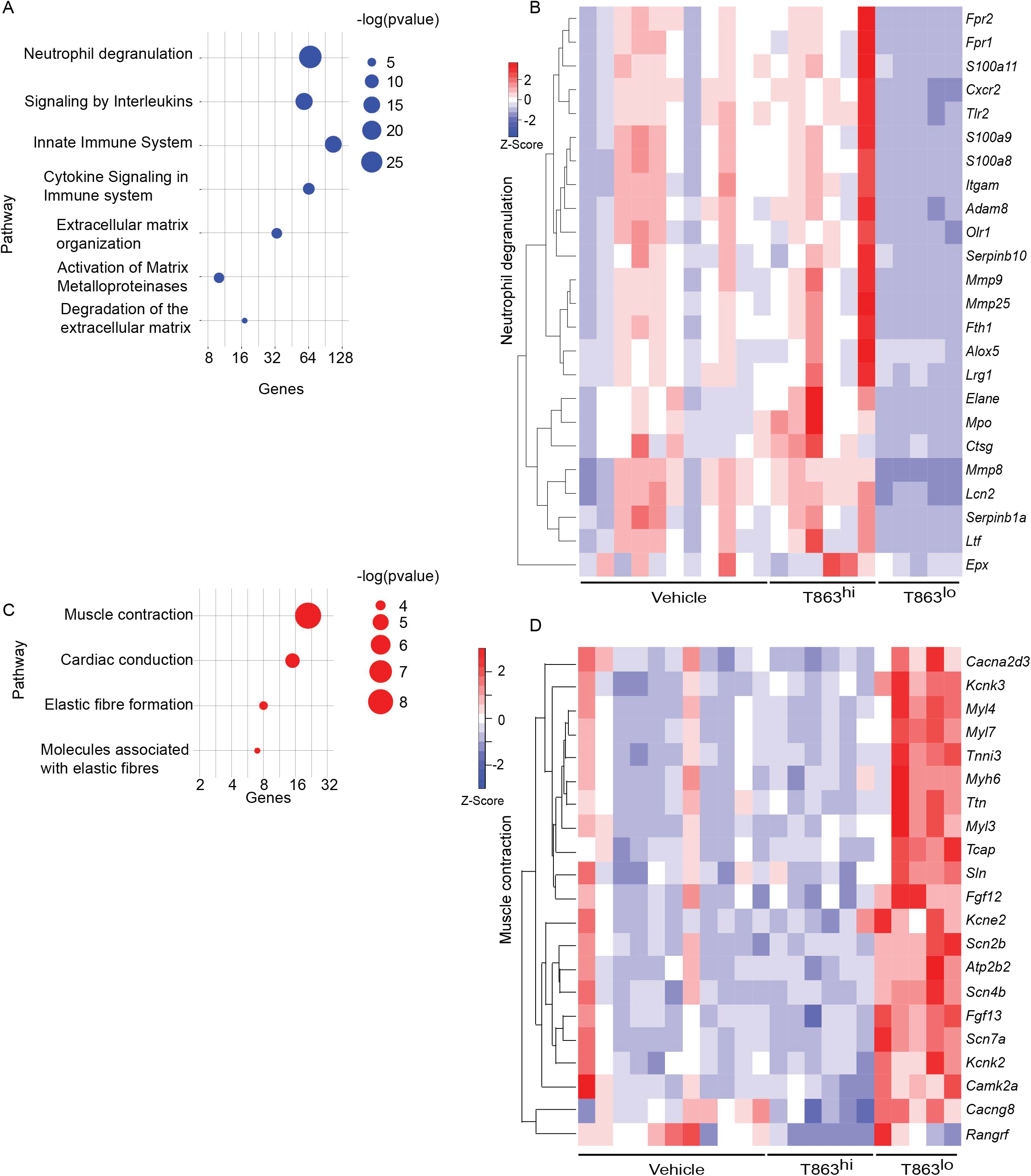
Correlates of DGAT1 inhibition mediated bacterial control from RNA-seq analysis. (A) Pathways negatively enriched in T863^lo^ group compared to T863^hi^. (B) Heat map of neutrophil degranulation associated genes that were differentially expressed between T863^lo^ and T863^hi^ or Vehicle groups. (C) Pathways positively enriched in T863^lo^ group. (D) Heat map of muscle contraction associated genes that were differentially expressed between T863^lo^ and T863^hi^ or Vehicle groups.

CD11b, a neutrophil cell surface marker, is an essential integrin required for neutrophil recruitment to sites of inflammation. S100A8 and S100A9, abundant cytosolic markers of neutrophils form a heterodimer that are important for the expression of CD11b on neutrophils and recruitment of neutrophils to TB granulomas (Scott et al., 2020). A reduced expression of *S100a8* and *S100a9* in T863^lo^ animals compared to T863^hi^ and vehicle group was a strong indicator of reduced neutrophil recruitment. Neutrophil recruitment also relies on CXCR2 engagement with CXCL1 on its cell surface. CXCL1 can be made by several cell types including neutrophils but is essential for neutrophil recruitment. Importantly CXCL1 was identified as the most prominent determinant of TB susceptibility across outbred strains of mice (Niazi et al., 2015). Reduced expression of both *Cxcl1* and *Cxcr2* suggested reduced neutrophil recruitment in the T863^lo^ animals (Fig 3B and Table S1). Myeloperoxidase (MPO), a marker for azurophilic granules that is secreted upon neutrophil activation, was used to verify if the abundance of neutrophils is indeed altered upon T863 treatment for 2 weeks. In five out of six vehicle treated animals, intense staining for MPO was found clustered in granulomatous regions (Fig 4A). Such staining was evident in only three out of six T863 treated animals (Fig 4A and B). To understand whether bacterial numbers correlated with the extent of MPO staining, we stained serial lung sections with SYBR Gold using acid fast staining methods. While variable degree of acid fast staining was evident in the vehicle treated group, it was negligible in the T863 treated animals that exhibited lower neutrophil recruitment (Fig 4C and D). MPO staining correlated with acid fast staining, with 3 out of 6 T863 treated animals exhibiting negligible MPO and SYBR Gold staining (p=0.0036) (Fig 4E). Together, these data indicate that T863 dependent lowering of bacterial burden is associated with reduced neutrophil recruitment and activation.

**Figure 4.**
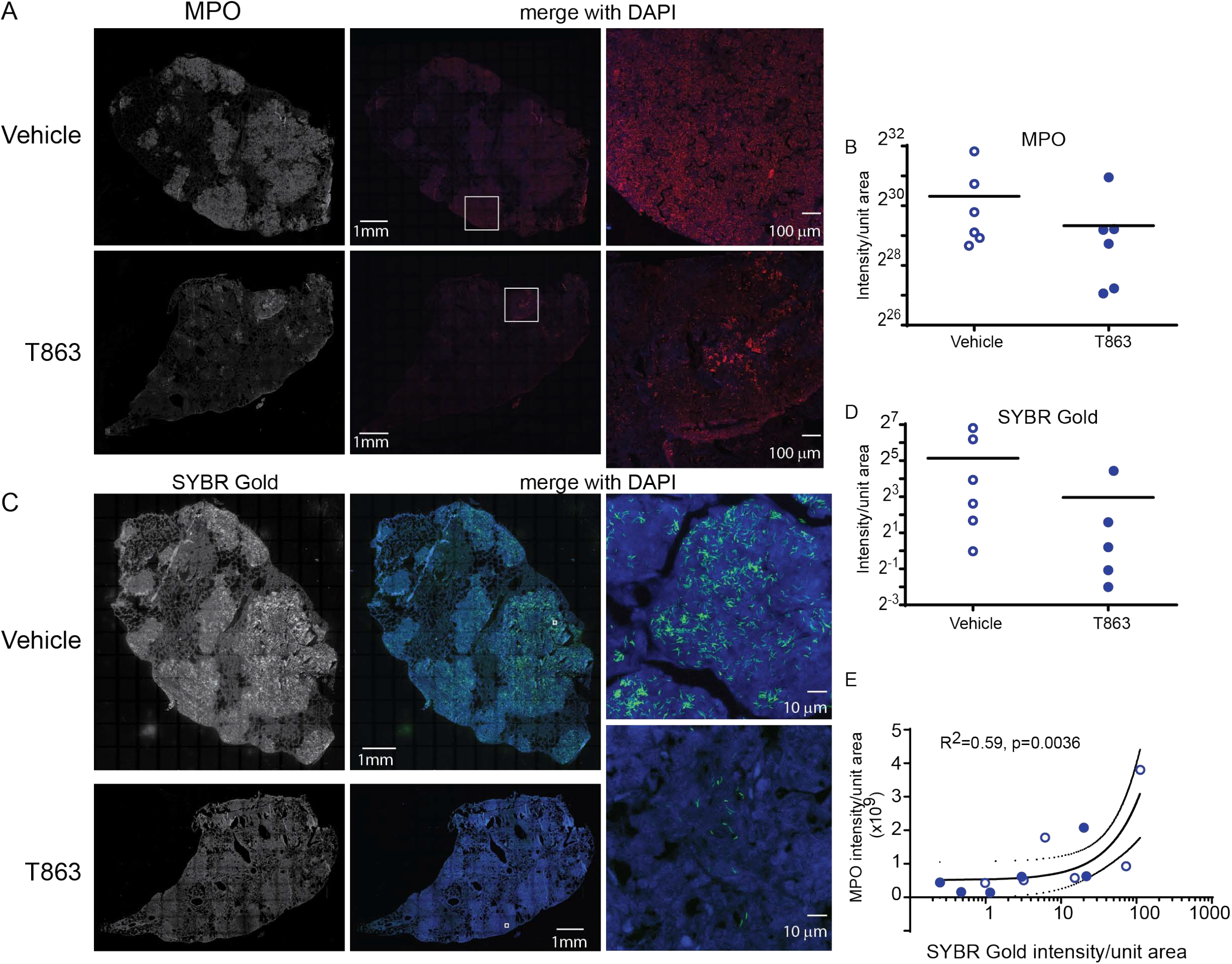
MPO levels correlate with reduced bacterial burden in T863 treated mice. Immunostaining (A) and quantification (B) for MPO in lung tissue sections from indicated groups of mice. The left panel is a raw intensity image of MPO stained whole lung section stitched together. The middle panel is the psuedocolored MPO signal (red) merged with that for DAPI (blue). Panel on the right is a conofcal image on an indivdual granuloma from the same lung section acquired at 10X magnification from indicated regions of the middle panel (white box). (C) Acid fast staining using SYBR Gold in same lungs as in (A). The middle panel is the pseudocolored SYBR Gold signal (green) merged with that for DAPI (blue). Panels on the right are images at 100X magnification from indicated regions from the middle panel (white box). (D) Quantification of SYBR Gold signal. Data in (B) and (D) are from 6 animals per group with horizontal bar representing the mean of each group. (E) Pearson’s correlation of MPO signals and SYBR Gold signals from (B) and (C). x-axis is log10 transformed only for visualization purposes.

### DGAT1 inhibition alters the pro-inflammatory cytokine profile

Previous studies have demonstrated that in both human and mouse macrophages, triglyceride synthesis by the enzyme DGAT1 enhances the pro-inflammatory response to both *M. tuberculosis* and LPS (Castoldi et al., 2020; Jaisinghani et al., 2018). However, the inflammatory responses *in vivo* in a granulomatous disease could be the sum total of contribution from various cell types and signals. Pathway analysis of the RNA-seq data also suggested reduced expression of several cytokines and genes involved in cytokine signalling in the T863^lo^ group (Fig 3A). The *sst1* locus of C3HeB/FeJ mice confers susceptibility to tuberculosis which has recently been shown to be driven by expression of IL1Ra, a secreted IL1 receptor antagonist which is part of the Type I IFN response (Ji et al., 2019). We first evaluated the kinetics of the pro-inflammatory response in a high dose infection of C3HeB/FeJ mice. We observed a temporal increase in expression of the Type I IFN response genes *Il1rn* (Fig 5A), *Ifit2* (Fig 5B), *Ifit1* and *Ifih1* (data not shown) upon infection. The kinetics of pro-inflammatory genes such as *Tnfα*, *Il1β*, and *Il6* were not identical to that of the type I IFN response, with peak induction of *Il6* observed at d14 (Fig 5C), and that of *Tnfα* observed at d28 (Fig 5D), while *il1β* expression remained constantly raised to similar levels across these time points (Fig 5E). This high dose of infection thus correlated well with the previously described kinetics of the inflammatory response (Bhattacharya et al., 2020; Ji et al., 2019) and was detectable during sustained neutral lipid accumulation.

**Figure 5.**
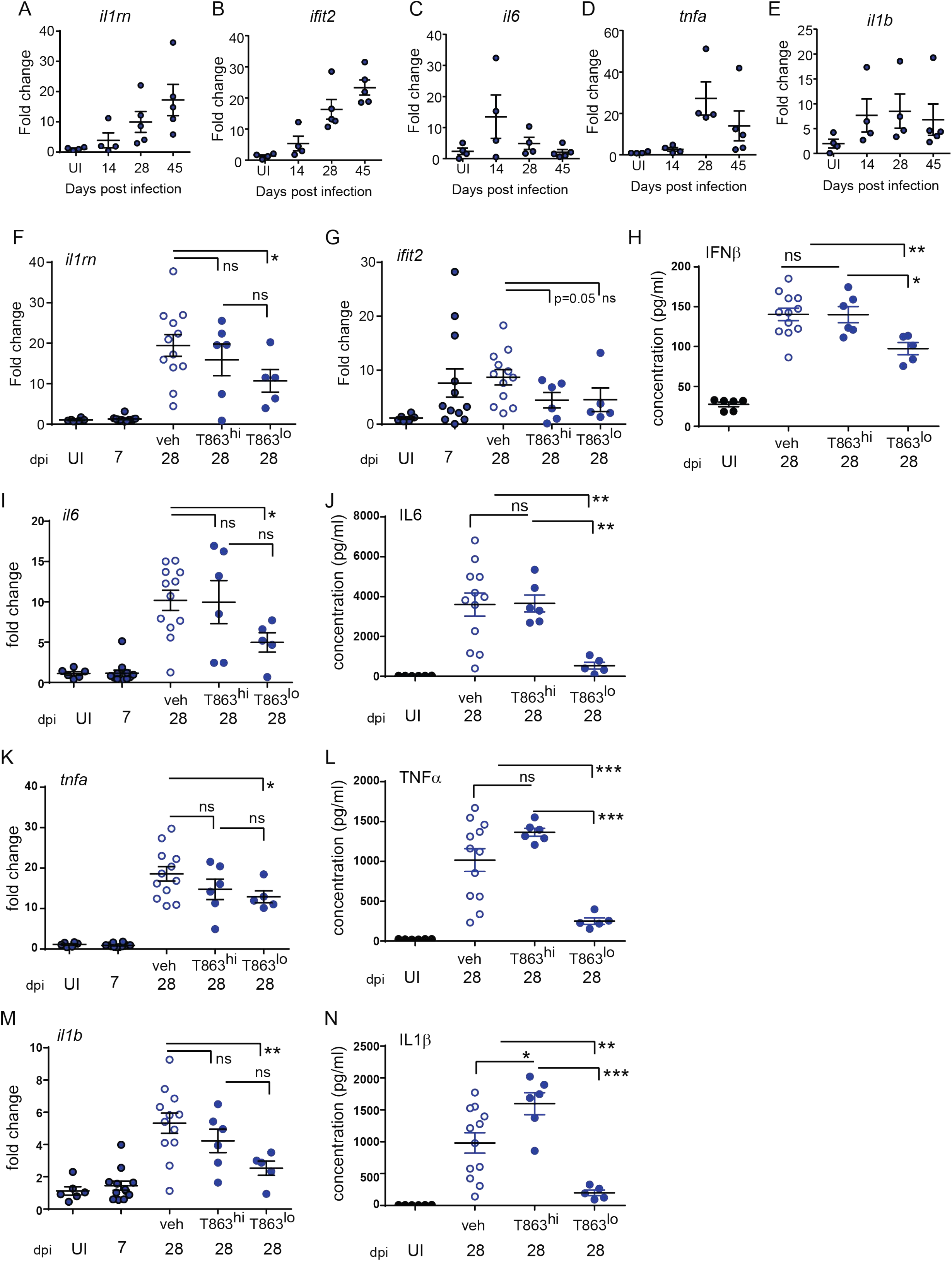
Progression of Type I IFN and pro-inflammatory cytokine response is limited by DGAT1 inhibition in animals exhibiting bacterial control. (A-E) Quantitative real time PCR based gene expression of indicated genes in uninfected (UI) mice and indicated days post infection. Data are normalized to mean of uninfected group. qRT-PCR (F,G,I,K,M) and multiplex ELISA (H,J,L,N) based estimation of gene expression and protein levels in lungs of induicated groups of mice. qRT-PCR data are normalized to mean of uninfected group. T863^hi^ refers to animals that had bacterial burden similar to the vehicle group and T863^lo^ refers to animals that exhibited less than 25% bacterial burden of the vehicle group. Horizontal bars in each dataset represent mean+/- SEM. Statistical analysis:unpaired t-test with Welch’s correction; ***p<0.001, **p<0.01, *p<0.05, ns=not significant.

The induction of *Dgat1* as early as d14 post infection led us to question whether DGAT1 activity in the granuloma affects the sustained inflammatory response to infection. We evaluated whether granuloma TG levels contribute to the local inflammatory cytokine expression, and whether this correlated with bacterial control. *Il1rn* expression was ~2 fold lower in T863^lo^ compared to the vehicle group (Fig 5F). Although the T863^hi^ group exhibited a decrease, this was not statistically significant. Expression of *Ifit2* was also two folds lower in the T863 treated group, albeit independent of bacterial control (Fig 5G). IFNβ being the upstream cytokine of the Type I IFN response, we measured lung levels of this cytokine. IFNβ levels were found to be elevated in the lungs at d28 post infection in the vehicle group compared to uninfected animals. Lungs of T863^lo^ animals harboured 1.4 fold lower levels of IFNβ (Fig 5H) while no significant difference was found in the levels of IFNβ between the T863^hi^ group and vehicle group. Transcript abundance of *Il6* and levels of IL6 protein were also found to be 2-fold and 7-fold lower in the T863^lo^ group compared to the vehicle group while they were not altered in the T863^hi^ group (Fig 5I and J). Transcript abundance of *Tnfα* was reduced 1.4-fold and the cytokine levels were reduced 4-fold in T863^lo^ group compared to vehicle group (Fig 5K and L). Similarly, transcript abundance of *Il1β* was reduced 2-fold and the cytokine levels were reduced 4-fold in T863^lo^ group compared to vehicle group (Fig 5M and N). Together with the RNA-seq results, these data confirm that decrease in bacterial burden upon T863 treatment was associated with decreased pro-inflammatory response *in vivo*.

### T863 alters eicosanoid flux during Mtb infection

Triglycerides have been proposed as a source of lipid mediators of inflammation known as the eicosanoids (Dichlberger et al., 2014). Eicosanoids are oxidation products of arachidonic acid that function in an autocrine and paracrine manner to regulate processes such as vasodilation, infiltration, cytokine production, and tissue healing. The relative abundance of prostaglandin E2 to leukotriene A4 is a well-established determinant of bacterial control in the C57BL6 mouse model (Mayer-Barber and Sher, 2015). Additionally, the zebrafish and mouse model of mycobacterial infection also indicates that optimal ratios of LXA4 and LTB4 are essential for control of infection (Bafica et al., 2005; Tobin et al., 2010). The role played by several other eicosanoids remains unclear. To assess if these lipid mediators of inflammation are altered as a result of T863 treatment, we measured eicosanoids using multiple reaction monitoring based mass spectrometry in lung tissues. We detected several oxidation products from each of the cyclooxygenase, Cyp450 oxidase, or lipoxygenase dependent pathways. Cyclooxygenase dependent prostaglandins PGE2 and PGJ2 were found to be slightly lower in T863 treated animals although the decrease was not statistically significant (Fig 6A,B). 11-HETE, another metabolite from the COX pathway was instead found to be elevated 1.48 fold (Fig 6C). Similarly, Cyp450 dependent 5,6-EET was also found to be elevated 1.74 fold (Fig 6D). LTB4 is a major determinant of disease susceptibility in the zebrafish model of mycobacterial granulomatous disease, with an imbalance in LTB4 to LXA4 levels driving disease susceptibility in both directions (Tobin et al., 2010). However, LTB4 was not altered in response to T863 treatment (Fig 6E). However, 5-HETE, another 5-LOX dependent metabolite was found to be 1.55 fold elevated in lungs of T863 treated animals (Fig 6F). Alox5/12-LOX dependent 12-HETE and Alox5/Alox15 dependent LXA4 were also found to be elevated 1.55 fold in these animals (Fig 6G and H). Interestingly, T863^hi^ subgroup had slightly higher levels of 5,6-EET, 5-HETE, and 12-HETE than the T863^lo^ subgroup of animals suggesting the presence of modifiers of the effect of T863 *in vivo*.

**Figure 6.**
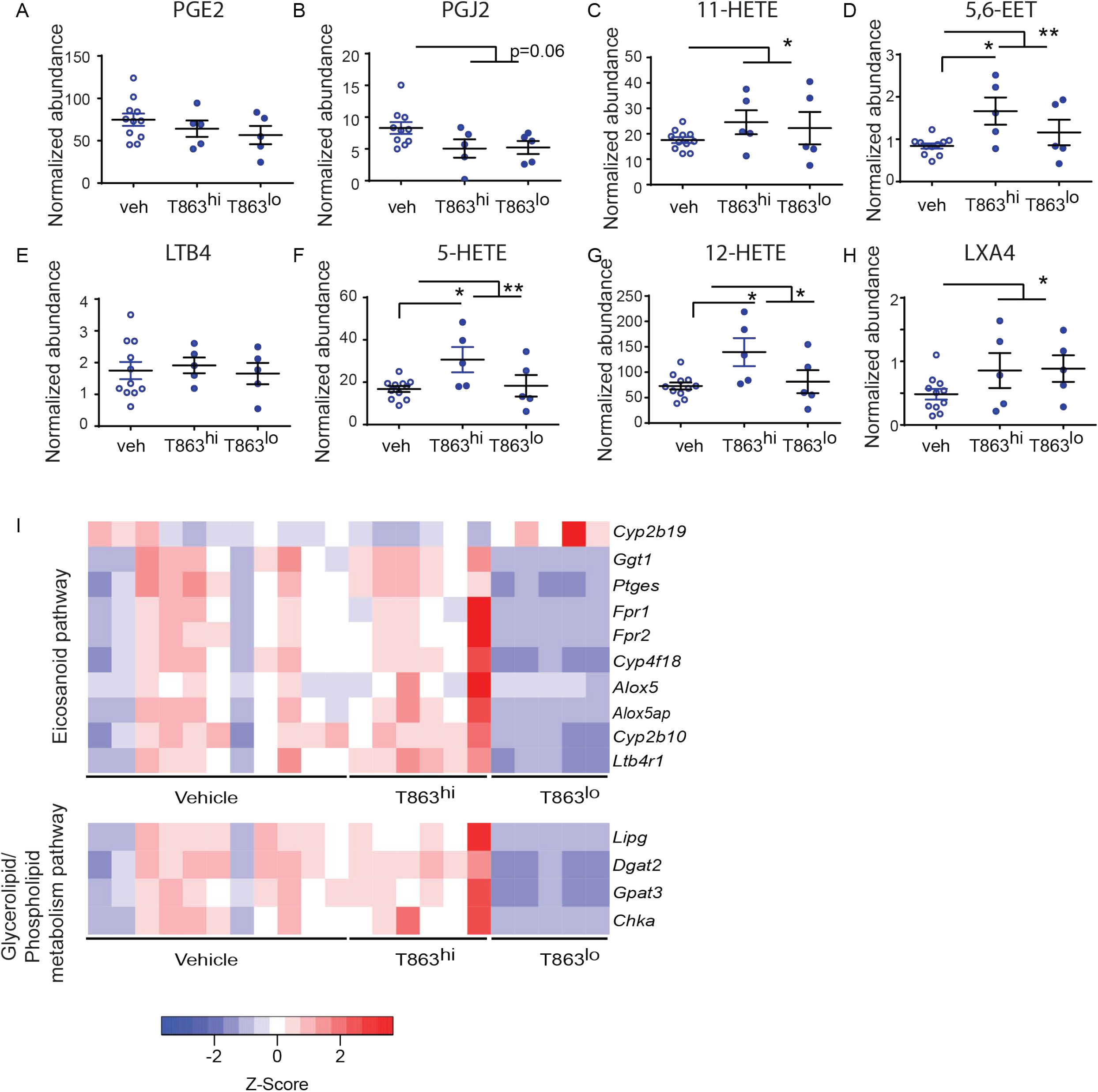
Dgat1 inhibition leads to alterd flux through the eicosanoid pathway. Arachidonic acid metabolites from the cyclo-oxygenase (COX) (A-C), cytochromeP450 (D), 5-lipoxygenase (E-F), 12-lipoxygenase (G), and 15-lipoxygenase (H) pathway that were measured by multiple eraction monitoring are indicated. Abundance for each metabolite has been normalized to an internal spike-in standard of the same class. Horizontal bars in each dataset represent mean+/- SEM. Statistical analysis: unpaired t-test with Welch’s correction. Vehicle group is compared to the combined T863 group and to each sub group. Only statistically significant differences are indicated by asterix, All other comparisons are not statistically significant **p<0.01, *p<0.05. (I) Heatmap of differentially expressed genes belonging to eicosanoid synthesis and sensing and glycerolipid/phospholipid metabolism pathway.

We then asked if genes involved in eicosanoid metabolism and sensing are potential modifiers of eicosanoid levels that distinguish the T863^hi^ and T863^lo^ groups. We manually curated a list of genes known to be involved in eicosanoid production and response. We compared expression of these genes across T863^hi^, T863^lo^ and vehicle treated animals. This analysis revealed that expression of *Ptges* was downregulated in T863^lo^ animals compared to all other groups (Fig 6I). We also found decreased expression of *Alox5* and *Alox5ap* (ALOX-5 activating protein) in the T863^lo^ group compared to both Vehicle and T863^hi^ group. These genes are required for conversion of arachidonic acid to 5-HETE which is then converted to 12-HETE by 12-lipoxygenase and LXA4 by 15-lipoxygenase. This finding is consistent with reduced levels of 5-HETE and 12-HETE in the T863^lo^ group compared to T863^hi^ group. Similarly, lower expression of the putative arachidonic acid hydroxylase *Cyp4f18* and arachidonic epoxygenase *Cyp2b10* in lungs of T863^lo^ group was consistent with lower levels of 5,6-EET in T863^lo^ group. Fpr1/Fpr2, a neutrophil specific receptor of several agonists including LXA4, was found to be lower in the T863^lo^ group. No other lipoxin receptor was found to be affected in expression. Expression of the LTB4 receptor *Ltb4r1* was also reduced statistically significantly in T863^lo^ animals (Fig 6I). Given that LTB4 was not altered at the metabolite level, the overall effect in T863^lo^ animals would have been dampened signalling from the LTB4 axis compared to the vehicle group. Our data reveal that *in vivo*, not all arachidonic acid metabolites correlate with neutral lipid levels and there seems to be a more intricate balance and regulated flux of arachidonic acid selectively across these pathways, and DGAT1 activity is indeed a major regulator of this flux. Arachidonic acid, a requisite precursor of eicosanoids, is a fatty acid chain present in glycerolipids and phospholipids. We found lower expression of the endothelial lipase *Lipg* and of genes involved in cellular glycerolipid and phospholipid synthesis (*Dgat2*, glycerol-3-phosphate acyltransferase 3-*Gpat3*, choline kinase alpha-*Chka*) in the T863^lo^ group (Fig 6I). Together, these data point towards diminished flux towards neutral lipid, phospholipid, and lipoxygenase dependent eicosanoids to be associated with T863 mediated control of bacterial burden.

## Discussion

Triglycerides have been considered inert storage form of excess fatty acids which can provide energy to power the cell’s energy demands. In case of TB infection, triglyceride rich foamy macrophages are conspicuously present at the cellular cuff of necrotic granulomas. Studying their relevance *in vivo* requires use of a model that exhibits these pathological findings. C3HeB/FeJ mice, similar to guinea pigs and humans, exhibit this feature of TB infection. Diacylglycerol-O-acyltransferases are required for the conversion of diacylglycerol to triacylglycerol. Here, we demonstrate that DGAT1 is not only induced upon infection, but is also essential for TG accumulation in the granuloma. In this study we found that DGAT1 inhibition can lead to better control of bacterial growth *in vivo*, via reducing the detrimental inflammatory immune response to infection.

The pro-inflammatory response in tuberculosis must be balanced at a critical threshold to achieve bacterial control and prevent tissue damage. Chronic inflammation and excessive production of TNFα cytokine is associated with higher risk of developing severe TB disease pathology (Roca et al., 2019; Tobin et al., 2010). In addition increase in Type I IFN response in Mtb infection is linked with host susceptibility to the disease (Ji et al., 2019), severe lung inflammation and neutrophil mediated NETosis (Moreira-Teixeira et al., 2020). IL1Ra, a Type I IFN induced protein that inhibits IL1β signalling, if deleted rescues disease severity in TB susceptible mice (Ji et al., 2019). T863 treated animals that exhibited decrease in bacterial burden, not only displayed significant reduction in both the pro-inflammatory response and Type I IFN response at the level of these genes, but also TNFα and IFNβ at the protein level, leading to a functional reduction in neutrophil recruitment. *Mycobacterium tuberculosis* induces neutrophil extracellular trap (NET) formation; though these NETs trap the bacteria, they fail to kill Mtb (Ramos-Kichik et al., 2009). Excess neutrophil recruitment (Scott et al., 2020) and neutrophil mediated NETosis have been shown to play a detrimental role in the host (Moreira-Teixeira et al., 2020). A neutrophil driven Type I IFN gene signature in the blood of TB patients, which declines upon antibiotic treatment, further provides support to the link between neutrophil driven pathology and bacterial growth (Berry et al., 2010). Our data identifies local triglyceride synthesis within the granuloma to be a driver for this axis.

While Mtb is able to utilize triglyceride from host macrophages, it is not an essential carbon source for the bacilli (Jaisinghani et al., 2018; Knight et al., 2018). Macrophage polarization to M1 versus M2 types creates distinct metabolic states *in vivo*, which impose distinct metabolic states of the intracellular bacteria (Huang et al., 2018). However, we found that neither M1 nor M2 polarization of murine macrophages rendered intracellular Mtb dependent on host TG synthesis. These data along with the effects on neutrophil recruitment pointed towards an alternate mechanism of why T863 could reduce bacterial burden. We thus questioned whether other possible products of triglycerides could alter the *in vivo* infection outcome. Eicosanoids are possible oxidation products of fatty acids derived from triglycerides (Dvorak et al., 1983). Indeed, our study revealed that lowering triglyceride levels led to lowering of prostanoid levels while it led to increase in levels of lipoxygenase-5 derived eicosanoids. Importantly, eicosanoid production and action seems to be regulated at the transcript level upon inhibiting triglyceride synthesis. This is in agreement with previous studies in murine bone marrow derived macrophages where inhibition of DGAT1 in M1 polarized macrophages led to decrease in levels of prostaglandins via expression of the gene *ptges* (Castoldi et al., 2020). Previously, prostaglandin production by IFNγ polarized, Mtb infected BMDMs was also shown to be partially triglyceride dependent (Knight et al., 2018). Our data identifies additional effects of triglyceride lowering on eicosanoid production *in vivo*, some of which occur through metabolite flux and some through transcriptional control. Alox5 is critical to the production of several eicosanoids such as 5-HETE, 12-HETE, LTB4 and LXA4. Previous studies on *alox5*^-/-^ mice have identified a pro-bacterial role of this gene in TB infection (Bafica et al., 2005). Our finding of reduced *alox5* expression associated with decrease in bacteria burden is consistent with this. Bafica et al., demonstrated that the reduction of bacterial burden in *alox5*^-/-^ mice could be reversed by administration of a stable LXA4 analog. Our data suggests that there may be additional lipoxins that may play similar roles in the C3HeB/FeJ mouse model as lowering of granuloma triglyceride levels led to increase in LXA4 levels despite reduced levels of *alox5* in T863^lo^ animals while increase in 5-HETE and 12-HETE only in T863^hi^ animals and not in T863^lo^ animals. The notable impact of triglyceride lowering on bacterial burden, seen only in a subset of mice, could be due to simultaneously lower levels of 5,6-EET, 12-HETE, and 5-HETE. 12-HETE and 5-HETE are both drivers of neutrophil infiltration and their increase in the T863^hi^ group relative to the vehicle group may be driving neutrophil mediated inflammation which prevents decline in bacterial burden (Mishra et al., 2017). A combination of gene and metabolite level control in T863 treated animals therefore leads to suppression of neutrophil mediated inflammation. Further investigations on the role of these metabolites in driving inflammation and neutrophil driven disease pathology will be important in addressing these questions. The impact of altering specific eicosanoids during an active infection may be a useful strategy to enhance host directed control of infection. This idea is supported by previous studies in Mtb infected C3HeB/FeJ mice where treatment with low dose aspirin or Ibuprofen, inhibitors of PGE2 production and enhancers of lipoxin production, led to suppression of mycobacterial growth and promoted resolution of inflammation (Kroesen et al., 2018; Vilaplana et al., 2013).

Our study provides proof-of-concept whereby targeting neutral lipids plays a protective role in TB progression. DGAT1 inhibitors have been a focus of pharma investment, primarily as a means to treat obesity and metabolic effects of diet induced obesity (DeVita and Pinto, 2013). The involvement of DGAT1 in dietary lipid clearance from enterocytes and maintenance of gut health have also encouraged development of parenteral routes of delivery of DGAT1 inhibitors (Denison et al., 2014; Haas et al., 2012; Kumar, 2017). Consistent with this, intravenous administration of the DGAT1 inhibitor in our study prevented lowering of serum TG with no effect on animal weight but led to lowering of granuloma TG levels. Given the need to deliver anti-TB therapy in large affected populations in resource limiting settings, tissue specific DGAT1 targeting would be the way forward. Decreased expression of *Gpat3*, *Dgat2*, and *Chka* in animals that decreased bacterial numbers in response to T863 suggests a potentially additive effect of decreasing neutral lipid storage towards limiting the detrimental immune response. Moreover, inhibition of triglyceride synthesis potentially alters functions of lipid droplet localized proteins. Lipid droplet localized proteins that regulate innate immune defense might be useful additional targets for developing host directed therapies against TB.

## Materials and Methods

### Mice

C3HeB/FeJ mice (stock 000658; The Jackson Laboratory) were purchased from Jackson laboratory; mice were maintained and bred in-house in accordance with the CSIR-Institute of Genomics and Integrative Biology’s animal ethics committee’s guidelines and approval. Mice were 6-10 weeks old when used. Equal numbers of male and female mice were used for each infection experiment. During the course of maintenance and infection, mice were fed standard chow diet and water *ad libitum* and kept under standard 12 hours dark and light cycle.

### Bacterial cultures and mice infection

Frozen stocks of Erdman strain of *Mycobacterium tuberculosis* were thawed and cultured in Middlebrook 7H9 broth supplemented with Albumin-Dextrose-Saline. Mid log phase cultures were used for infections. Mice were infected with Erdman strain of *Mycobacterium tuberculosis* using inhalation exposure system, Glas Col, LLC at 100 cfu per animal (low dose infection) and 500 cfu per animal (high dose infection). All infections were performed with actively growing, early log phase cultures.

### DGAT1 Inhibitor (T863) administration

A selective and potent DGAT1 inhibitor T863 (Sigma-SML0539) was used in this study. Mice were injected intravenously via tail vein with 100 μl of 633.76 μM T863 solution or the vehicle control (1X PBS, 0.05% tween-80, 20% ethanol). To achieve solubilization of T863 for intravenous dosing, 4 mg of T863 was weighed and dissolved in 800 μl of ethanol to which 10 μl of 20% Tween-80 was added. This suspension was sonicated in a water bath sonicator, followed by addition of 800 μl ethanol and 10 μl of 20% Tween80, followed by water bath sonication. After solubilization, 1.6 ml of 10X PBS and 11.2 ml of tissue culture grade water was added to achieve a solution of 250μg/ml. This would achieve approximately 17μg/ml per mouse. Treatment was initiated either at day 7 or day-14 post infection with intravenous dosing every three days till day-28 post infection.

For oral dosing of T863, mice were orally administered T863 at 30mg/kg, at 5μl/g of body weight. The animals were dosed for 15 consecutive days starting at day-14 post infection and at day-28 post infection mice were given T863 and vehicle dose in the morning and then sacrificed later that day. The dose concentration were adopted from (Cao et al., 2011). Compound was dissolved/ solubilized in 0.1% carboxymethylcellulose (CMC): 0.1% of Tween-80, required amount of the compound T863 was weighed and transferred to a mortar pestle and required equal volume of 0.1% carboxymethylcellulose (CMC): 0.1% of Tween-80 was added, it was ground well and transferred to a tube, vortexed and sonicated in a water bath sonicator. Dissolved compound was vortexed and sonicated prior to each dosing schedule. The usability of these vehicles for oral administration of a test compound was shown by (Singh et al., 2012).

### Histopathological analysis of lung samples

Left caudal lung lobe was harvested in a vial containing 10% neutral-buffered formalin and kept at room temperature overnight for fixation, lung lobes were soaked in gradient of sucrose solution 20% in 1X PBS overnight and then in 30% sucrose solution overnight at 4°C prior to cryo-sectioning at a section depth of 5μm using *Leica Cryostat CM 1850*. Sections were stained with Hematoxylin and Eosin as described previously in (Jaisinghani et al., 2018).

### Histology for small intestine

Small intestine from the orally administered T863 and vehicle group of mice, which were infected with low dose of Mtb (100CFU/animal) were carefully harvested. Small intestine were stretched out gently in a petri plate and flushed gently with the 10% neutral-buffered formalin solution to clean out the internal contents as well as fix the intestinal tissue. Prior to cryo-sectioning of the intestinal tissue, fixed small intestine was stretched out and cut into three equal parts: proximal, mid and distal, cryo-sectioning of the intestinal tissue was done using an improved swiss-rolling technique described previously by (Bialkowska et al., 2016). Section were further used for studying the neutral lipid clearance in small intestine by staining with Oil red O and hematoxylin.

### Hematoxylin and Oil Red O Staining

The sections were first dipped in a pre-warmed Oil Red O (Cat no. 09755, Sigma) solution (0.18% in 60% isopropanol) for 1 h at 60°C, and then rinsed twice with water for 5 min each. The slides were then dipped in hematoxylin solution for 5 min, rinsed, and mounted in 90% glycerol. Images were acquired using Leedz Microimaging 5 MP camera attached to a Nikon Ti-U microscope. For quantifications of the signal, whole lung sections were scanned using a 1800dpi document scanner and red channel intensity quantified using Fiji. The Lab color space module was chosen to select thresholds.

### Lung and spleen cfu analysis

Lung lobes were harvested in 7 mL vial containing 2 mL of PBST (0.05% Tween-80) and 1 mm glass beads, lung and spleen tissues were homogenized using a homogenizer and the procedure followed for homogenizing was to give pulse for 2-3 minutes in 30 seconds pulse duration, 100 μl of the lung homogenates from each sample was taken for CFU analysis and rest of the homogenates were stored in different tubes in −80°C for lipid extraction and cytokine measurement. Dilutions of the neat lung and spleen homogenates were prepared and plated in 7H11 agar (BD Biosciences, USA) with 10% oleic acid-albumin-dextrose-catalase (OADC, Himedia laboratories, India). Colonies were counted 20-21 days later and calculated.

### BMDM isolation and infection

BMDMs were isolated from femurs and tibias of male C3HeB/FeJ mouse, femurs and tibias were excised carefully and put in a RPMI media, extra tissues attached to the bones were removed, head of the bones were cut and gently flushed the bone marrow cells in the media, bone marrow cells were homogenized and passed through a 0.7μm filter and centrifuged at 1500 rpm for 5 minutes. Cells were suspended in 10 ml of RBC lysis buffer and incubated for 5 minutes; cells were resuspended in RPMI media containing 20% L929 cells supernatant and differentiated in non-tissue culture treated petri plates, cells were incubated at 37°c under 5% CO_2_. On 7^th^ day media were removed from the petri plate and 4 mL 1x PBS + 0.5mM EDTA were added to each petri plates and incubated for 3-4 minutes, cells were harvested in a falcon and centrifuged at 1500 rpm, cell pellet was resuspended in 30 mL RPMI media, cells were counted and seeded. BMDMs were seeded in a 48-well plate in RPMI with 10% FBS media, cells were pretreated with mouse recombinant IFN-γ (BMS326, Invitrogen) at 6.25ng/mL for overnight and followed by infection with *Mycobacterium tuberculosis*, Erdman strain at MOI-1, after 3 hours of phagocytosis, extracellular bacteria were removed by washing with 1xPBS for four times and then fresh RPMI with 10% FBS media were added along with same concentration of IFN-γ, 10μM of T863 (Sigma-SML0539) and DMSO control were added in designated wells after 3 hours of infection. CFU plating were done at 3hours, Day-2, Day-4 and Day-6 post infection, during the experiment, at day-2 and day-4 media were changed and IFN-γ, T863 and DMSO were added along with the media in designated wells.

Similar experiment protocol was followed for IL-6 stimulation; BMDMs were pretreated with IL-6 (200-06, Peprotech) at 10ng/mL for overnight and infection protocol was followed as above. For M2 polarization, cells were pretreated with recombinant IL-13 (PMC0134, Thermo Scientific) at 20ng/mL and IL-4 (PMC0045, Thermo Scientific) at 20ng/mL concentration overnight and infection protocol was followed as above and T863 and DMSO treatment were also followed as describe above.

### Global transcriptome profile and differential expression analysis

A modified NEBNext RNA Ultra II directional protocol was used to prepare the libraries for total RNA sequencing. The first step involved the removal of ribosomal RNA of cytoplasmic and mitochondrial origin using biotinylated, target-specific oligos combined with rRNA removal beads. Following purification, the RNA was fragmented using divalent cations under elevated temperature. Next, the cDNA was synthesized using Reverse transcriptase and random hexamers in a first strand synthesis reaction. Subsequently, the cDNA was converted to double stranded cDNA where Uracil was added instead of Thymine. The strand specificity was preserved by a USER enzyme based digestion of second strand thereby leaving one functional strand which maps to the DNA strand which it was transcribed from. The USER digested single strand molecules were enriched and indexed in a limited cycle PCR followed by AMPure bead purification to create final cDNA library for sequencing. Sequencing reads from Illumina HiSeq X were checked for quality using FastQC and fastq files were trimmed and adapter sequences were removed using Trimmomatic (Bolger et al., 2014) and were aligned to the mm10 mouse reference genome using HISAT2 (Kim et al., 2019). We performed post alignment processing with SAMtools (Li et al., 2009) to put the alignment data into a format suitable for downstream analysis. To quantitate the expression levels of genes, we generated a gene read count matrix using feature Counts (Liao et al., 2014). Normalized counts and differentially expressed genes were obtained using DESeq2. Statistical tests for differential expression were performed with the DESeq2 package (Love MI et al., 2014) of the Bioconductor suite in R. For final results, we set the log2 FC >1 and <-1 cut off of DEGs for up-regulated and down-regulated genes respectively and p-values of < 0.05. Further ToppGene suite was used for gene list enrichment analysis.

### Immunofluoroscence of mice lung tissue

lung tissues were processed for cryosectioning as described above in the histopathological analysis method section, lung tissue sections of 5μM thickness were rehydrated in MilliQ water in a Coplin jar for 10 minutes. Sections were dip in 1x TBS in a coplin jar for 10 minutes followed by permeabilization in 0.25% tween-20 in 1x TBS for 10 minutes. Sections were rinsed thrice with 1x TBS for 10 minutes each. Tissue sections were then incubated with blocking buffer (10% donkey serum + 1% BSA in 1x TBS) at room temperature for 1 hour. Primary antibody (Myeloperoxidase, MPO Cat No AF3667) were added at a conc. of 5μg/mL and incubated for overnight at 4°C in a humidified chamber. After primary antibody incubation sections were taken out of 4°C and kept at room temperature for 20 minutes and followed by washing thrice with 0.025% triton-x100 in 1x TBS in a coplin jar for 10 minutes each. Sections were incubated with secondary antibody at 1:200 dilutions (Donkey Anti-Goat IgG NL557, Cat No NL001) for 1 hour at room temperature, sections were rinsed thrice with 1x TBS for 10 minutes each and then stained with DAPI (cat no. 10236276001, Roche) for half an hour at room temperature. Sections were washed thrice with 1x TBS for 5 minutes each, sections were air dried and mounted with anti-fade diamond mountant (Cat no. P36961, Invitrogen). Mounted samples were kept in dark to let it dry and then observed and imaged using Leica SP8 confocal microscope. Signal intensity per unit area was quantified using Fiji software.

### SYBR Gold Acid-fast staining of *Mycobacterium tuberculosis* on mice lung tissue

SYBR Gold staining of *Mycobacterium tuberculosis* was shown as an improved and efficient method to stain M. tuberculosis than the routinely used acid-fast staining methods Ziehl Nelson and fluorescent acid-fast staining methods like Auramine Rhodamine and Auramine-O (Ryan et al., 2014). Tissue sections were taken out from −20°C and kept at room temperature for 20 minutes and sections were rehydrated in MilliQ water for 10 minutes. Briefly, SYBR gold stain (cat no S11494, Invitrogen) was diluted 1:1000 in a staining solution (phenol crystals 8g, glycerine 60mL, Isopropanol 14mL and distilled water 26mL). Slides were kept on a dry bath set at 65°C and SYBR Gold staining solution was dropped gently on the tissue sections and kept it for 5 minutes. After 5 minutes slides were removed from the dry bath and let it cool for a minute, sections were washed with acid alcohol (0.5% HCl, 70% Isopropanol) for 3 minutes. Sections were further washed with water for 10 minutes. After washing tissue sections were stained with DAPI at 1:1000 dilutions (2mg/mL stock solution) for 30 minutes and then washed with 1x PBS 3 times of 5 minutes each. Sections were mounted with prolong anti-fade diamond mounting medium and then observed and imaged using Leica SP8 confocal microscope.

### Lung RNA isolation and qRT-PCR analysis

Lung lobes taken for RNA isolation were stored in RNA Later solution (AM7021, Invitrogen) in −80 freezer, lung tissues were thawed on ice, after thawing, tissues were put in 2mL screw cap vials containing 1mm glass beads and 1mL of RNAzol RT solution for homogenization using a rabbit homogenizer and the procedure for tissue homogenization was to give 30 seconds pulse and 30 seconds rest on ice for a total of 3 minutes, total RNA isolation was carried out following the recommended specifications of RNAzol RT Sigma, aqueous phase separation and RNA precipitation both were carried out in BSL3, after the precipitation, samples were removed from BSL3 and further steps were carried out in the molecular biology lab. RNA samples were further purified using the RNeasy mini kit (Cat No 74106, Qiagen) and later equal amount of RNA samples were treated with Turbo DNAse, cDNAs were prepared from 1μg of RNA using Verso cDNA synthesis kit (Cat. No AB1453B, Thermo Fisher Scientific). Quantitative real time PCR was performed on Light cycler 480 (Roche) instrument with Kapa Sybr Fast Universal qPCR master mix (Cat. No SFLCKB, Roche). Transcripts levels were normalized to a housekeeping gene 18s rRNA and following primers were used in this study:

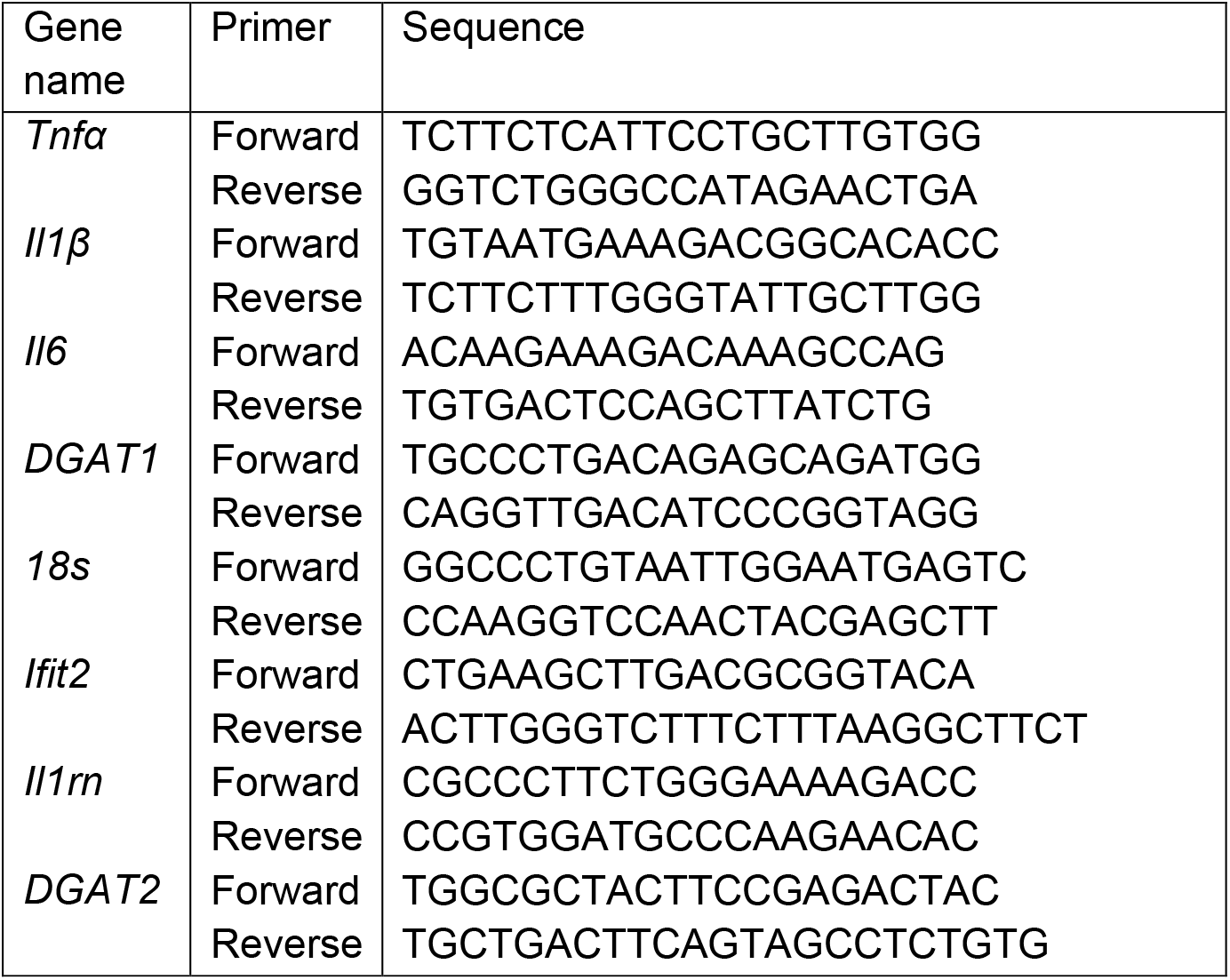

### Cytokine measurements

An aliquot of lung homogenates were stored for cytokine measurements, protease inhibitor cocktail was added to the homogenates and it was centrifuged at low speed at 3000rpm for 5 minutes to settle the tissue debris down, supernatant was transferred to a new tube and stored in −80°C. Lung homogenates stored for cytokine measurement were thawed on ice, samples were centrifuged at high speed at 12,000 rpm for 10 min, supernatant was transferred to a new tube, and samples were filtered through 0.2μm spin-cup filter to remove all Mtb present in the sample. Filtered cell free lung homogenate were then taken out of BSL3 and used for cytokine measurement, cytokines were measured using the recommended protocol from ProcartaPlex Multiplex Immunoassay (Cat no. PPX-10-MX2W9GN, Invitrogen) and data was acquired on MAGPIX, Luminex system.

### Serum triglycerides quantification

Serum triglyceride levels were measured using Triglycerides-250T Cobas Integra 400+ biochemistry analyzer reagent (Cat No.20767107322) on COBAS INTEGRA 400 plus system.

### Extraction and analysis of Eicosanoids

Eicosanoids extraction protocol was adopted from (Zhang et al., 2015). Eicosanoids from mouse lung were extracted by liquid-liquid extraction. Briefly, lung homogenates were prepared as described above, 200μL of lung tissue homogenates from each samples were added to 500 μL of methanol (2% formic acid) spiked with internal standard mixture (10ng for each) and mixed vigorously by vortexing for 5 minutes. Tubes were centrifuged at 12,000 g for 10 min at 4°C, after centrifugation the supernatant was transferred to a new tube. Water (700μL) and Ethyl acetate (500μL) were added to the supernatant. The sample was mixed vigorously for 2 min and centrifuged at 12000 g for 10 min. The upper organic phase was transferred to a new tube and the aqueous phase was extracted again by adding Water (700μL) and Ethyl acetate (500μL).

Organic phase was combined and 1 mL of chloroform was added in the end, followed by evaporation of the solvent completely. Each sample was re-suspended in 300μL of 100% ethanol and further filtered through a 0.2 micron nylon syringe filter (cat no SLGNX13NL) prior to injection into the UPLC system. Eicosanoids were chromatographically separated using an Exicon LC system with a Waters ACQUITY UPLC BEH C18 reverse phase column (2.1 x 50 mm, 1.7μm). The following binary gradient was used to resolve eicosanoids: Solvent A: Water + 10mM Ammonium Acetate; Solvent B: Acetonitrile. A 10 min run was performed at a flow rate of 0.5 mL/min with 5% solvent B upto 1min, 50% upto 2.5 min, 70% upto 3.5 min, 80% upto 6.0 min, 90% upto 7.0 min, 98% upto 7.5 min, then decreased to 50% upto 8.0 min, increased back to 80% upto 8.5 min, and maintained at 80% upto 9.5 min. Solvent B was reduced to 5% by 9.6 min and maintained until 10 min.

The runs were acquired through an AB SCIEX QTRAP 6500+ LC/MS/MS system in low ass range using Analyst 1.6.3 software. The Q1 and Q3 masses for the multiple reactions monitoring (MRM) based analyte detection are mentioned in the table below.

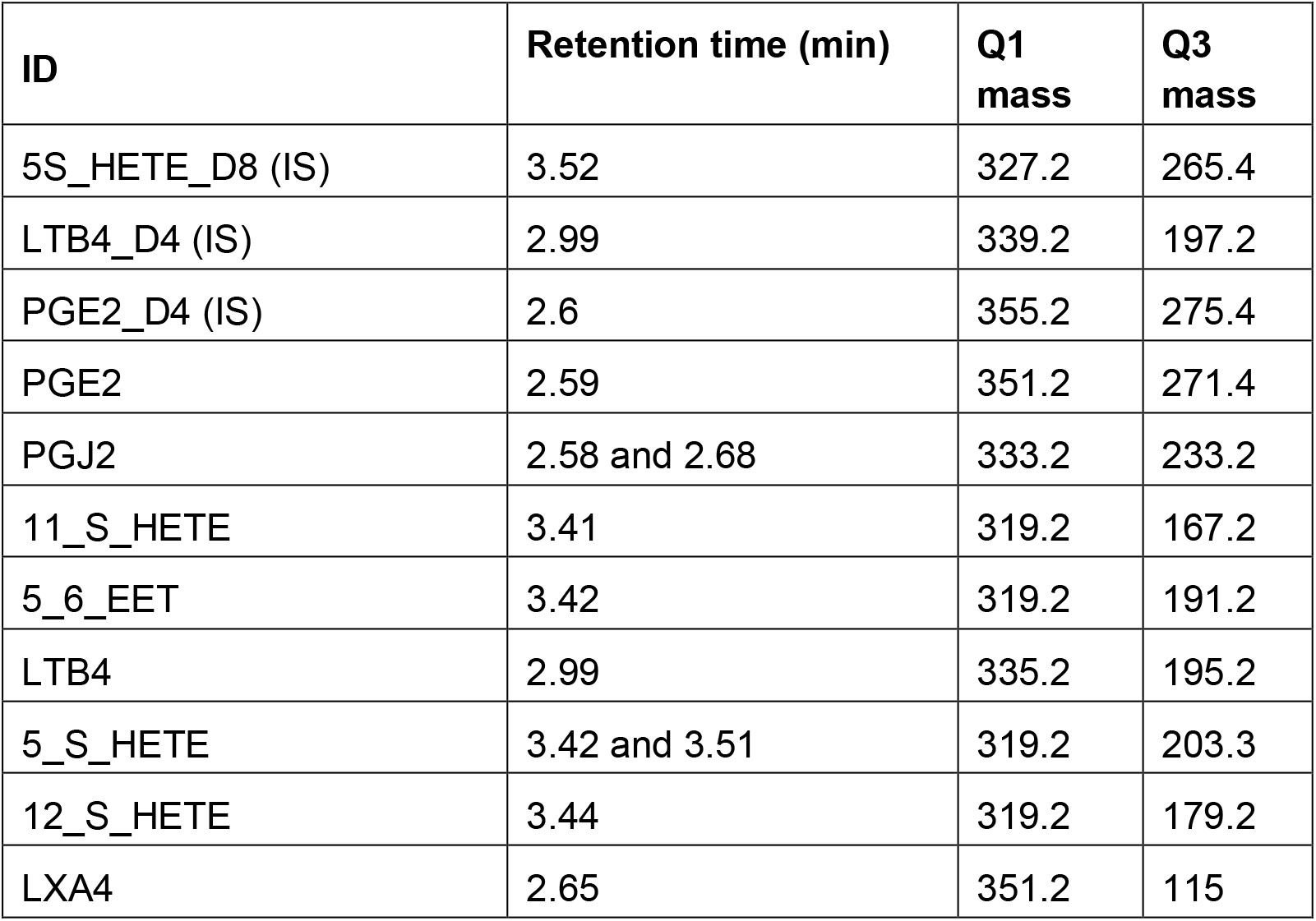

The ion detection parameters for each class of eicosanoids were further optimized for the in house mass spectrometer using commercially available deuteriated spike-in standards based on the technical note available from SCIEX. Signal intensities for each metabolite were normalized to the signal intensity of deuteriated spike-in standards in each sample belonging to the same class.

### Statistical analysis

All statistics were computed using Prism. T-test with Welch’s correction was used for comparison between selected groups. Pearson’s correlation was calculated between signal intensity/unit area for MPO and SYBR Gold for the full lung section per animal.

## Author contributions

Conceptulaization:SG; Study Design: SD, VR and SG; Data analysis: SD and SG; Animal experiments: SD, MM, SG, VR and PA; RNAseq analysis: PA and SD; Eicosanoid measurement: DM, AB, MM; Manuscript writing: SD and SG; Manuscript reviewing and editing: all authors; Funding procurement: SG

## Acknowledgements

The authors would like to thank Mr Digamber Prasad for BSL3 maintenance and Mr Santosh Singh, Mr Rajesh Yadav and Mr Amit Misri for animal husbandry, and Rajat Ujjainiya for help with serum triglyceride measurement. The authors also thank Dr Shantanu Sengupta for the mass spectrometry facility.

## Ethics Statement

The animals were housed and handled at the CSIR-Institute of Genomics and Integrative Biology Mathura Road Campus according to directives and guidelines of the Committee for the Purpose of Control and Supervision of Experiments on Animals (CPCSEA) as per ethics proposal #IGIB/IAEC/Oct2018/07.

---

**Figure S1.**
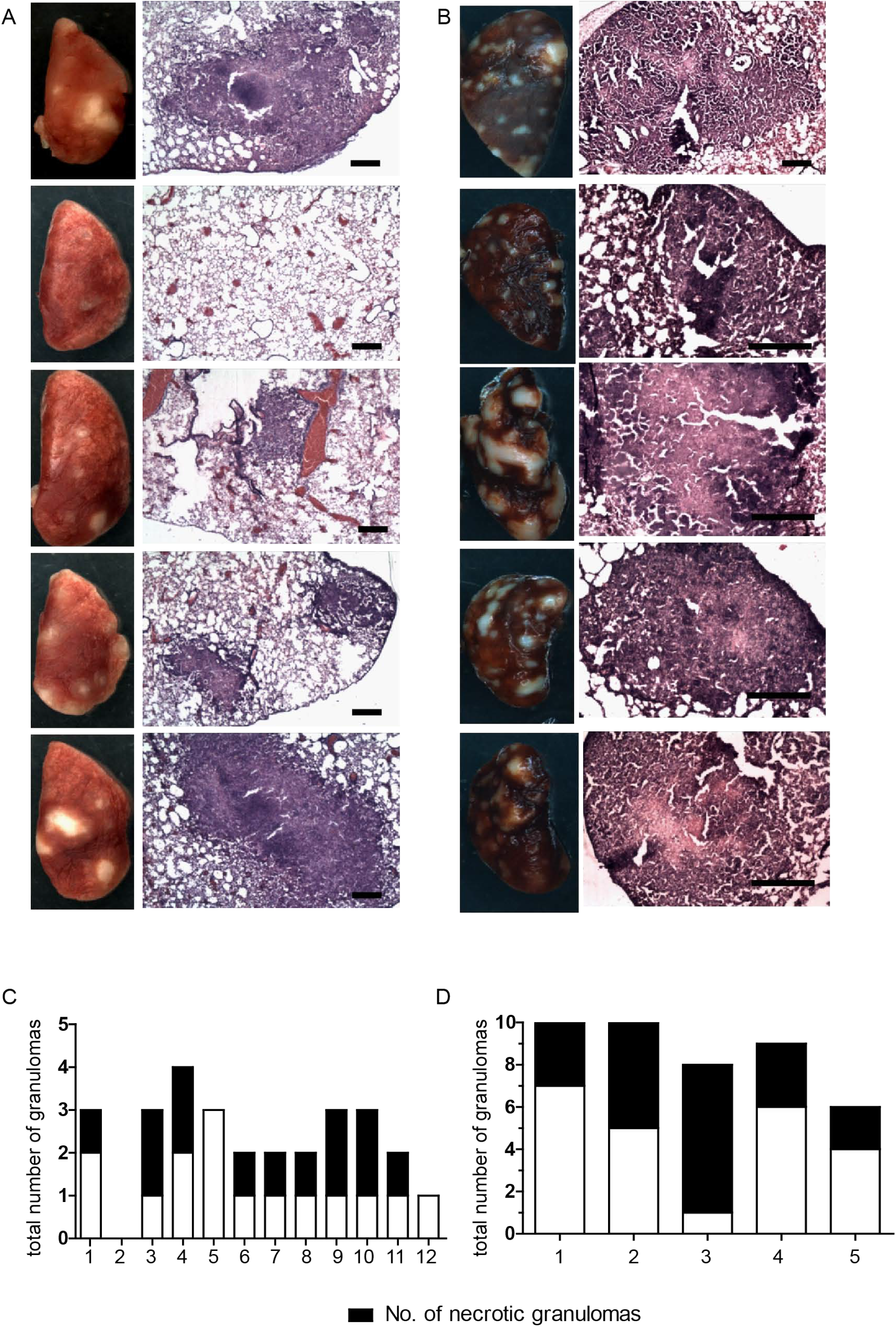
Gross pathology and cryosections from C3HeB/FeJ mice at d28 post infection with low dose (100 cfu) (A) and high dose (500 cfu) (B) of Erdman strain delivered via aerosol route. Section are stained with hematoxylin and eosin; scale bar=400 μm. Data are of 5 representative from low cfu and all 5 fromhigh CFU. Quantification of necrotic and total number of granulomas scored in the left caudal lobe of animals of low dose infection (C) and high dose of infection (D).

**Figure S2.**
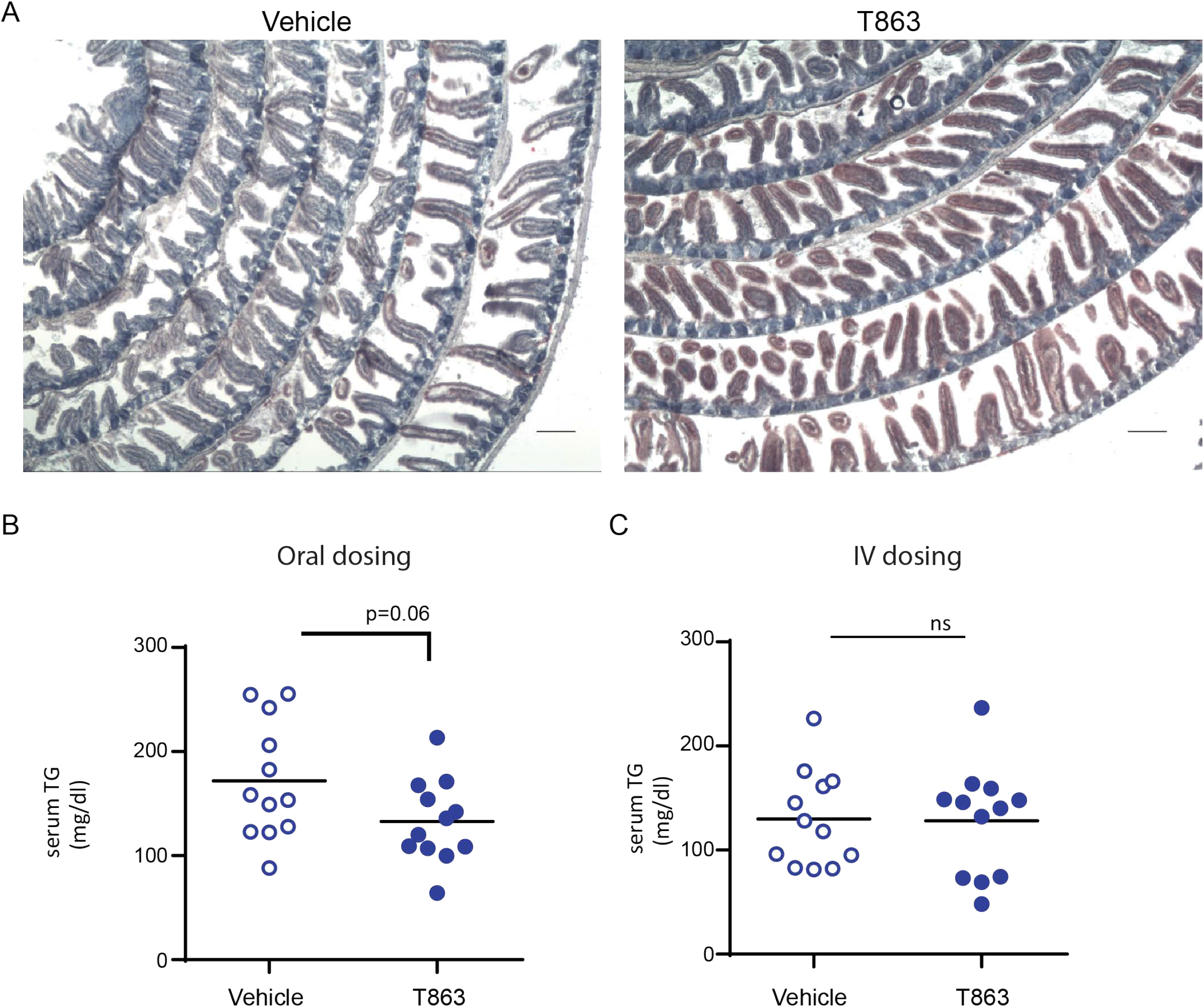
Effects of oral versus iv dosing of T863 on gut and serum TG levels. (A) Cyrosections of distal small intestine stained with oil red O and hematoxylin scale bar=200 μm. (B-C) Serum triglycerides in vehicle or T863 treated upon 3 weeks of oral (B) or intravenous (C) dosing.

**Figure S3.**
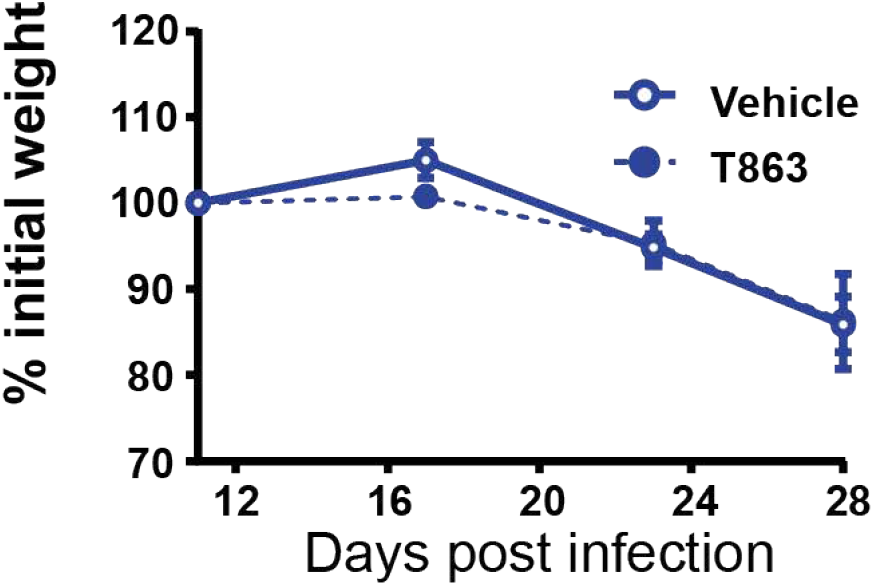
Effect of T863 treatment on weight loss over 3 weeks of treatment

**Figure S4.**
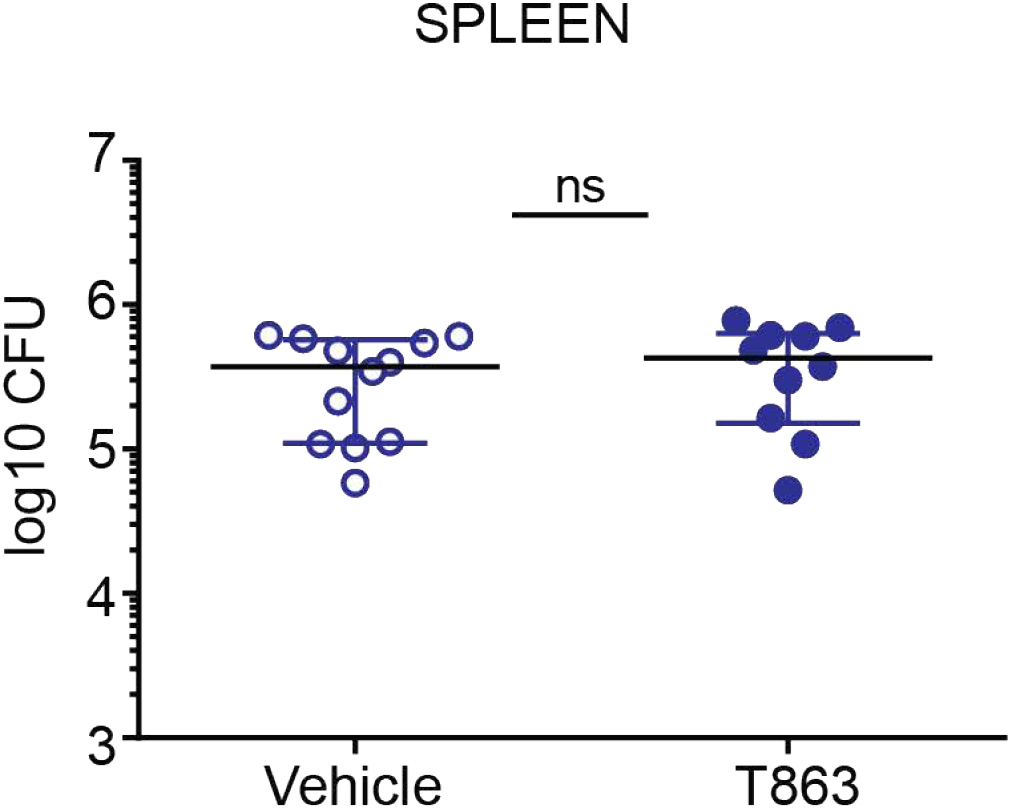
Splenic CFU at d28 from animals treated with Veicle or T863 for 3 weeks.

